# Characterizing Multimodal Mutational Processes in Cancer Using MOTET

**DOI:** 10.64898/2026.11.12.765471

**Authors:** Jiaqi Sun, Kelly L. Geyer, Natasha Q. Gurevich, Tao Guo, Masanao Yajima, Jonathan H. Huggins, Joshua D. Campbell

## Abstract

Mutational signatures are characteristic patterns of somatic alterations in cancer genomes. A single aberrant mechanism can simultaneously generate coordinated patterns of alterations across multiple mutation modalities, including single-base substitutions (SBSs), doublet-base substitutions (DBSs), and small insertions and deletions (IDs). However, these signatures are typically inferred separately and linked *post hoc*, obscuring their shared underlying mechanisms. To resolve this, we formally define a Mutational Process (MP) as a unified latent activity that induces coupled mutational signatures across these modalities. We developed MOTET, a Bayesian framework that jointly infers multimodal MPs. Applied to 2,583 whole genomes from the PCAWG cohort, MOTET recovers known MPs more accurately than standard post hoc correlation methods. MOTET also identifies a process linking an SBS37-related signature with a TG>CA-dominated DBS component and resolves canonical SBS12-like activity into two separate MPs with distinct tissue distributions. These results establish a paradigm for formally defining catalogs of MPs and demonstrate that integrating across modalities enhances the ability to discover new mutational patterns.

## INTRODUCTION

Mutational signature analysis has become a central framework for investigating the biological processes that shape cancer genomes^1–3^. Several signatures have been attributed to specific exogenous causes including tobacco smoking, ultraviolet (UV) radiation, or chemotherapy, as well as endogenous causes such as DNA-repair deficiency or aberrant APOBEC activity^4,5^. Determining the origins of these signatures could improve reconstruction of tumor evolutionary histories, identify previously unrecognized mutagenic exposures or repair defects, and inform cancer prevention and biomarker development^6,7^.

A single mutational process (MP) can generate coordinated patterns across multiple mutation modalities, including single-base substitutions (SBSs), doublet-base substitutions (DBSs) and small insertions and deletions (IDs). For example, tobacco smoking is associated with COSMIC signatures SBS4, DBS2 and ID3, whereas ultraviolet radiation is associated with SBS7, DBS1 and ID13^8,9^. These combinations provide complementary evidence of the underlying process and can help distinguish processes that are difficult to resolve using a single modality. Outside such well-characterized examples, however, the relationships among many SBS, DBS and ID signatures remain incompletely characterized.

Non-negative matrix factorization (NMF) methods have been widely used for de novo discovery of SBS, DBS and ID signatures^10–13^. In workflows that extract signatures separately within each modality, cross-modality relationships are subsequently inferred, often by correlating sample-level activities. Such associations can depend on cohort composition, sample size and correlation thresholds, and may connect mechanistically distinct processes that co-occur across tumors. Grouping can also be ambiguous when a signature from one modality correlates with several signatures from another. In PCAWG, different extraction and correlation procedures produced different networks of associated SBS, DBS and ID signatures^9^, and integration after extraction remains an approach used in recent large-scale studies^14^. Other methods jointly extract signatures through concatenated mutation catalogs or shared-exposure models, establishing the value of integrating information across mutation classes^9,15,16^.

Bayesian formulations of NMF have enabled data-driven selection of the number of signatures^17^, incorporation of prior knowledge and joint analysis across studies^18^, and improved robustness to model misspecification with uncertainty quantification^19^. Multimodal approaches have used integrated exposures to cluster patients^20^ or modeled correlations between modality-specific activities, including SNV and structural-variation signatures^21^. MOTIVE jointly infers SBS, DBS and ID signatures within a Bayesian Poisson NMF model, explicitly modeling covariance among modality-specific activities during inference (Geyer, K. L. et al. Joint estimation of signatures across mutation modalities using multi-modal non-negative matrix factorization. Manuscript in preparation). Although MOTIVE shares information across modalities during inference, it does not explicitly assign modality-specific signatures to shared MPs; these groupings must subsequently be inferred from the modeled associations.

Here, we introduce MOTET, a Bayesian hierarchical NMF model that extends MOTIVE by introducing a shared latent activity for each MP, linking its modality-specific signature profiles. Each process has an allocation of mutations across SBS, DBS and ID, estimated jointly with its signature profiles and activity in each tumor, making cross-modality grouping part of the model itself (**Fig. 1c**). We benchmarked MOTET against exposure-correlation approaches to evaluate its recovery of established cross-modality mutational processes. Applied to 2,583 whole genomes from 37 tumor types in PCAWG (ICGC/TCGA Pan-Cancer Analysis of Whole Genomes Consortium, 2020^22^), MOTET recovers smoking- and UV-associated signature combinations, resolves SBS12-like activity into two processes, and identifies a process coupling SBS37-like and DBS17-like signatures. These findings demonstrate the utility of MOTET’s process-level representation for refining mutational signatures, characterizing their relationships across modalities, and generating hypotheses about their underlying mechanisms.

**Fig. 1:**
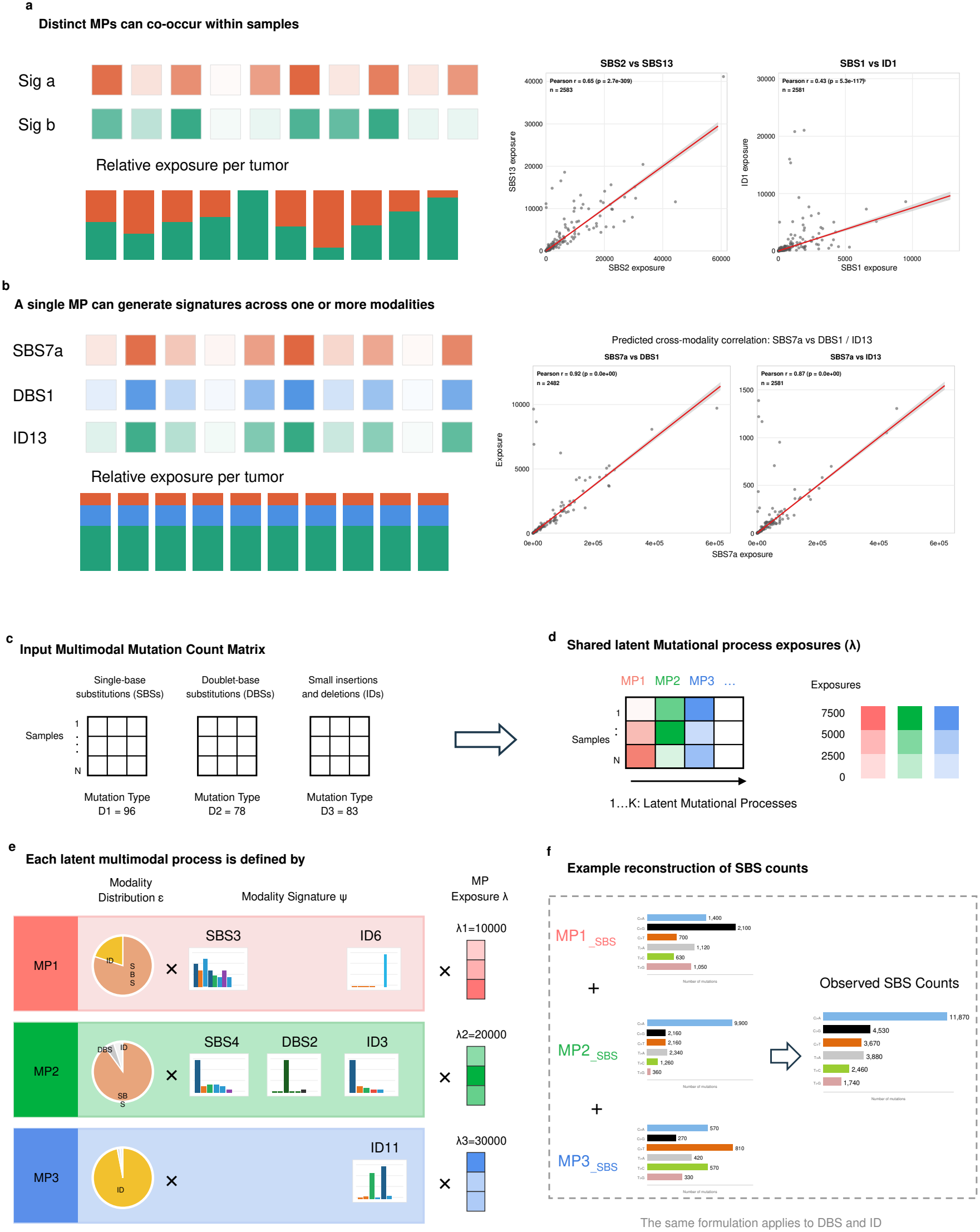
Conceptual framework of the MOTET model for joint extraction of multi-modal mutational signatures. **(a)** Distinct mutational processes (MPs) can co-occur within a tumor, and their relative contributions can vary across tumors. Each schematic column represents one tumor. Within each row, darker shading indicates greater exposure to that signature. Each stacked bar shows the fraction of the two signatures’ combined exposure attributable to each signature in that tumor. Scatterplots show separately estimated signature exposures in the Pan-Cancer Analysis of Whole Genomes (PCAWG) cohort for SBS2 versus SBS13 (n = 2,583 tumors) and SBS1 versus ID1 (n = 2,581). **(b)** A single MP can generate signatures in multiple mutation modalities. In MOTET, the expected exposures of its component signatures scale with process activity across tumors (shading), while their expected proportions remain fixed (stacked bars). Scatterplots show exposures for the ultraviolet radiation–associated signatures SBS7a versus DBS1 (n = 2,482) and SBS7a versus ID13 (n = 2,581) in PCAWG. In all scatterplots, each point represents one tumor. Red lines show linear regression fits, and gray bands indicate confidence intervals for the fitted mean. Pearson r and Spearman ρ are reported for each pair. **(c)** Input mutation count matrices for SBS, DBS and ID mutation modalities across the same set of N samples, with SBS96, DBS78 and ID83 classification schemes. **(d)** MOTET jointly infers a single exposure matrix λ, shared across the SBS, DBS and ID modalities, whose entries represent the sample-specific exposure of each K latent mutational processes (MPs). Color coding (pink, green, blue) denotes MP1, MP2 and MP3, respectively, and is maintained throughout b-d. **(e)** Each latent process is defined by three components: a sample-specific exposure λ, a modality distribution ε that specifies the relative contribution of the process to SBS, DBS and ID mutations, and modality-specific signature profiles ψ that define the mutation spectrum within each modality. **(f)** Each latent process contributes to the expected SBS counts through the product of its exposure, SBS modality weight, and SBS signature profile. These contributions are summed across processes to obtain the reconstructed SBS counts. The same reconstruction applies analogously to DBS and ID counts. Exposure values and count totals shown are illustrative and do not represent fitted output from a specific dataset; modality distributions and signature profiles are representative of the following biologically established processes: a homologous recombination deficiency-associated process (SBS3 and ID6), a tobacco smoking-associated process (SBS4, DBS2 and ID3), and an ID11-dominant process.

## RESULTS

### MOTET infers multimodal mutational processes

We define a mutational process (MP) as a latent source of mutations with a characteristic footprint in one or more of three mutation modalities: single-base substitutions (SBSs), doublet-base substitutions (DBSs), and small insertions and deletions (IDs). Distinct MPs can co-occur within tumors and vary in their relative contributions across tumors (**Fig. 1a**). In the Pan-Cancer Analysis of Whole Genomes (PCAWG^22^) cohort, SBS2 and SBS13 exposures estimated separately for each modality with musicatk^23^ were positively correlated (Pearson r = 0.65), yet several tumors had substantial SBS2 exposure with little estimated SBS13 exposure. SBS1 and ID1 exposures were also positively correlated (Pearson r = 0.43), although their relative contributions varied across tumors.

A single MP can also generate signatures in multiple modalities (**Fig. 1b**). MOTET models each process as contributing mutations across modalities in fixed expected proportions. As the activity of a process increases, the expected exposures of its component signatures therefore increase together. The ultraviolet radiation– associated signatures SBS7a, DBS1, and ID13^9,24^ illustrate this pattern. In PCAWG, SBS7a exposure was positively correlated with both DBS1 exposure (Pearson r = 0.92) and ID13 exposure (Pearson r = 0.87). Because exposure correlations alone cannot establish whether signatures share an underlying process, MOTET jointly models mutation counts across modalities to estimate each process’s footprint and its activity in each tumor.

MOTET takes matched count profiles for SBS with the 96 trinucleotide-context channels^2^, doublet-base substitutions (DBS) into 78 channels, and small insertions and deletions (ID) into 83 channels^9^ across the same set of samples as input (**Fig. 1c**) and infers up to *K* latent mutational processes represented by a single exposure matrix λ (**Fig. 1d**). Each latent process is defined by three components (**Fig. 1e**): its sample-specific exposure *λ*_*nk*_, a modality distribution *ϵ*_*k*_ that specifies the relative proportion of total SBS, DBS and ID mutations to the overall MP, a modality-specific signature profiles 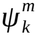 that describe the mutation pattern for each signature within each modality. The observed counts in each modality are reconstructed as the additive sum of contributions from all latent processes, illustrated here for the SBS path (**Fig. 1f**); the same formulation applies to DBS and ID counts, enabling coordinated mutational patterns across mutation modalities to be inferred directly within a single model.

In the example shown, the latent processes map to biologically interpretable combinations of signatures spanning one or more mutation modalities: a homologous recombination deficiency-associated process (SBS3 and ID6^6,25^), a tobacco smoking-associated process (SBS4, DBS2 and ID3^9,26^), and an ID11-dominant^9^ process. Together, these examples show how MOTET represents mutational processes across one, two or three mutation modalities, linking multimodal signatures through shared latent processes rather than post hoc exposure correlations.

### MOTET improves cross-modality process reconstruction relative to correlation-based approaches

We next evaluated whether MOTET improves reconstruction of cross-modality mutational processes relative to post hoc exposure correlation and established signature discovery tools^11,23^ (**Fig. 2a, b**). Correlation-based linking of separately extracted SBS, DBS and ID signatures is often difficult to interpret because a single SBS signature may correlate with multiple DBS or ID signatures, including signatures arising from distinct biological mechanisms or co-occurring processes in the same samples. We analyzed the combined pan-cancer whole-genome sequencing cohort from the Pan-Cancer Analysis of Whole Genomes (PCAWG) project (PCAWG_ALL; n = 2,583 samples; see Methods). In this cohort, post hoc exposure correlation produced a dense and threshold-sensitive network of putative SBS–DBS–ID associations (**Fig. 2c**). Using COSMIC^24^ version 3.6 annotations as the reference, the numbers of incorrect signature pairings and missed processes varied substantially with the correlation threshold (**Fig. 2h, i**), as did precision (**Fig. 2j**). MOTET results are shown as fixed benchmarks in these comparisons because its process assignments do not use a post hoc correlation threshold.

**Fig. 2:**
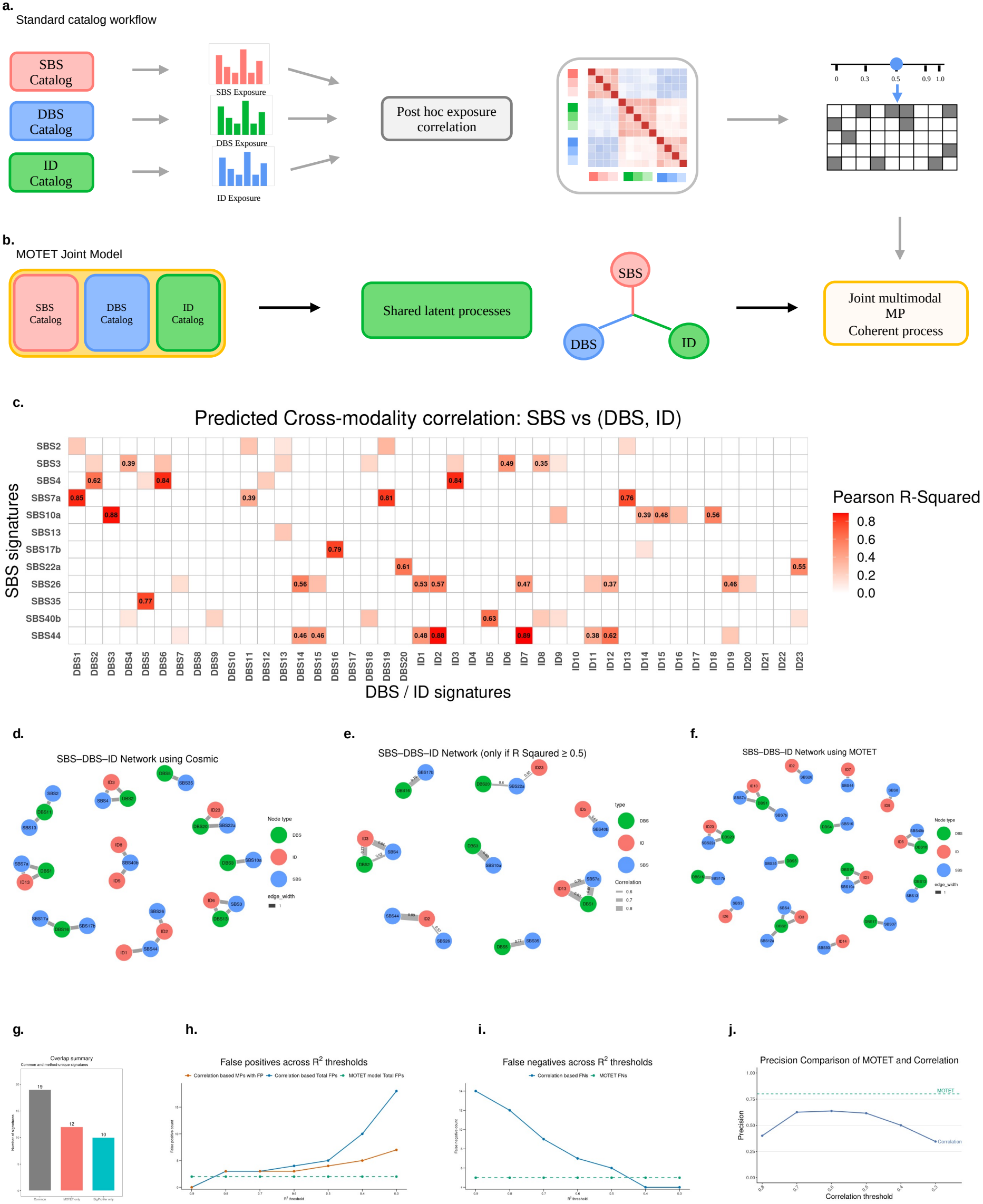
MOTET captures complex cross-modality mutational dependencies compared to standard catalogs. **(a)** Schematic of the standard catalog workflow: SBS, DBS and ID signatures are extracted and their exposures estimated independently for each modality, then linked post hoc using pairwise exposure correlations. Applying a correlation threshold defines groups of signatures with correlated exposures but can yield fragmented cross-modality links. (**b)** Schematic of the MOTET joint model: matched SBS96, DBS78 and ID83 count matrices are modeled jointly to infer shared latent processes directly, without a post hoc correlation threshold, yielding a joint multimodal mutational process with coherent cross-modality structure. **(c)** Matrix of squared Pearson correlation coefficients (r^2^) between the exposures of SBS and DBS/ID signatures independently inferred using musicatk^23^. The matrix is restricted to SBS signatures corresponding to the 12 COSMIC SBS signatures recovered by MOTET that have annotated DBS/ID associations in the COSMIC catalog (also shown in d). Colors and overlaid values indicate r^2^ for each SBS–DBS/ID pair. (**d)** Network of COSMIC-annotated cross-modality mutational processes involving the 12 SBS signatures shown in c and their associated DBS and ID signatures. Nodes represent signatures (SBS, blue; DBS, green; ID, red), and edges connect signatures annotated as belonging to the same process. **(e)** Network from d after retaining only edges with exposure correlations of r^2^ ≥ 0.5 in c and their connected nodes. Edge thickness is proportional to r^2^. **(f)** Network of 17 cross-modality mutational processes jointly inferred by MOTET. Node colors follow d; edges connect signatures within the same inferred process, and edge thickness carries no quantitative meaning. The network contains 15 connected components because two pairs of processes share a DBS signature: DBS1 links the SBS7b and SBS7a/ID13 processes, and DBS2 links the SBS12a and SBS4/ID3 processes. **(g)** Recovery of COSMIC SBS signatures from PCAWG_ALL by MOTET and SigProfiler, with 40 processes or signatures specified, respectively. Bars show signatures recovered by both tools (n = 19), MOTET only (n = 12), or SigProfiler only (n = 10). Recovery was defined as a cosine similarity ≥ 0.8 to a COSMIC SBS signature. **(h–j)** Comparison of the post hoc correlation approach across r^2^ thresholds (solid lines) with MOTET (dashed lines), using the COSMIC annotations in d as the reference. MOTET results remain constant because its process assignments do not use a post hoc correlation threshold. **(h)** Number of incorrect SBS–DBS/ID pairings (false positives). A pairing is classified as incorrect if the DBS or ID signature is not annotated as belonging to the same process as the SBS signature in the COSMIC reference. “MPs with FP” counts candidate processes containing at least one incorrect pairing; “Total FPs” counts all incorrect pairings across candidate processes. **(i)** Number of missed COSMIC-annotated processes (false negatives). A process is counted as missed if no candidate process recovers its annotated combination of SBS, DBS and/or ID signatures. **(j)** Precision, defined as the proportion of identified SBS– DBS/ID pairings supported by the reference: TP / (TP + FP).

MOTET recovered a larger set of COSMIC SBS signatures than SigProfiler^11^. The tools recovered 19 SBS signatures in common, with 12 additional signatures recovered only by MOTET and 10 recovered only by SigProfiler (**Fig. 2g**). We then examined cross-modality process recovery. The 12 SBS signatures with annotated DBS/ID associations corresponded to 10 distinct COSMIC-annotated processes (**Fig. 2d**). Post hoc exposure correlation at an R^2^ threshold of 0.5 recovered 8 of these 10 known processes (**Fig. 2e**). MOTET exactly recovered 8 of the 10 COSMIC-annotated processes. An additional inferred process contained the known SBS40b–ID5 pairing together with DBS18, a component not annotated for that process in COSMIC (Fig. **2f**).

Of the 17 cross-modality processes identified by MOTET, these 9 processes corresponded to known COSMIC processes, including the extended SBS40b–ID5–DBS18 process. Five others represented combinations absent from the COSMIC catalog used in this analysis, and the remaining 3 differed from their corresponding COSMIC-annotated compositions. We regard these 3 as uncertain rather than confirmed novel findings; for example, the SBS44–ID7^9,27^ association may reflect residual signal from the co-occurring ID2 signature rather than an independent process. Together, these results show that MOTET recovers established cross-modality processes and identifies additional candidate associations through joint inference, without requiring post hoc correlation-based linking.

### MOTET characterizes modality composition and exposure correlations among PCAWG mutational processes

We examined the modality composition of the 40 mutational processes inferred by MOTET in PCAWG_ALL (**Fig. 3a**). Of these, 34 were SBS-dominant, with SBS accounting for more than 50% of the modality distribution. Four processes either had an ID proportion above 50% or had a confident signature assignment only in the ID modality. None were DBS-dominant. The remaining two processes lacked a COSMIC signature match at cosine similarity ≥ 0.8 and did not meet our criteria for retaining candidate novel processes (see Methods). We therefore treated these as residual components rather than retained mutational processes. This distribution mirrors the broader composition of the PCAWG mutational landscape, where SBS signatures vastly outnumber DBS and ID signatures, and is consistent with the modality-specific sparsity priors used in the model^9^.

**Fig. 3:**
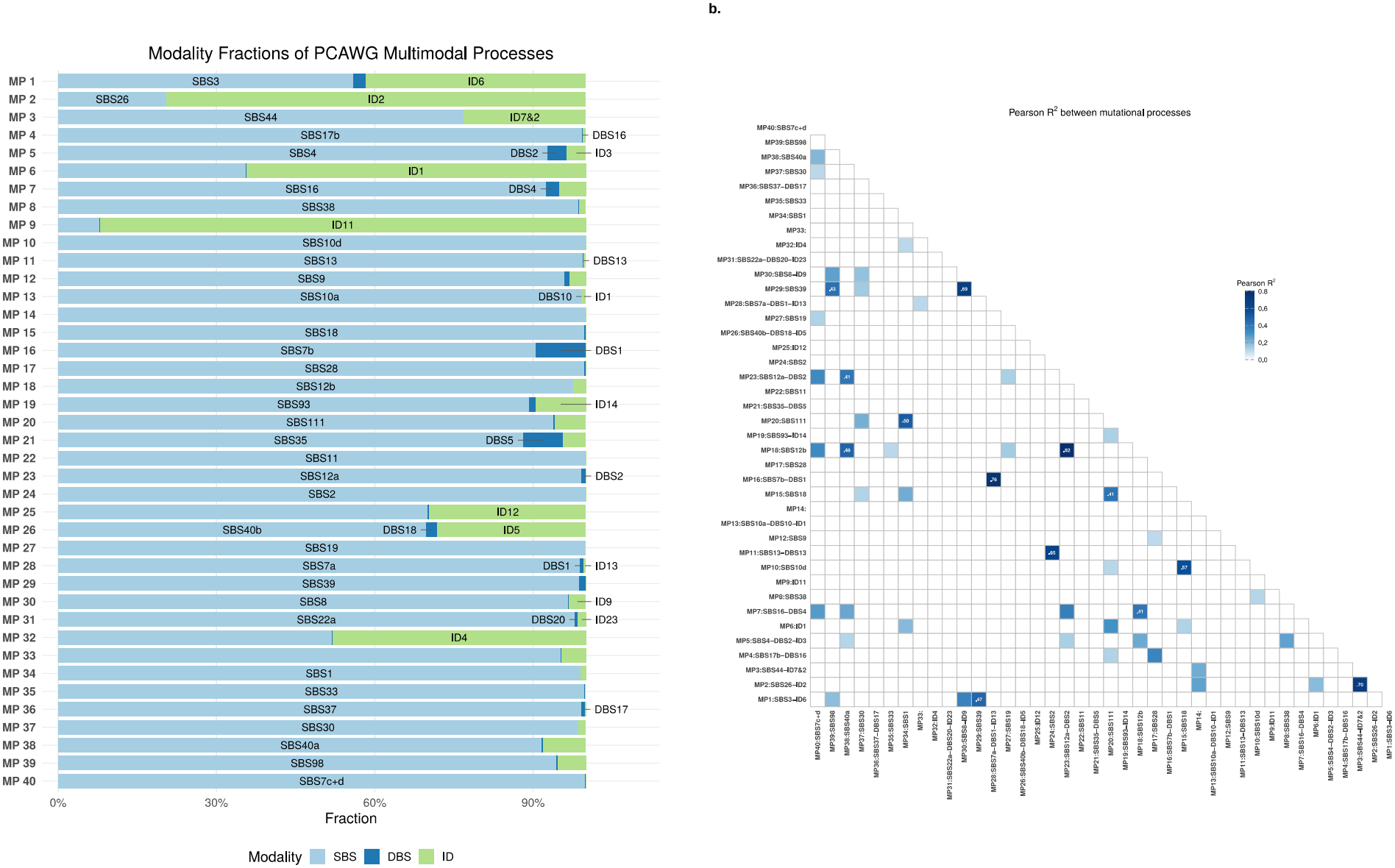
Modality distributions and correlations among composite mutational processes. **(a)** Modality proportions for the 40 mutational processes (MP1–MP40) inferred by MOTET in PCAWG_ALL (n = 2,583 samples; see Methods). Colors indicate single-base substitutions (SBS, light blue), doublet-base substitutions (DBS, dark blue) and insertions/deletions (ID, green). Proportions sum to 1 within each process. Labels identify the constituent signature(s) represented within each modality segment. **(b)** Squared Pearson correlations (r^2^) between per-sample exposures of the 40 mutational processes in PCAWG_ALL. Axis labels show MP numbers and their dominant constituent signatures. Only the lower triangle is displayed, excluding self-correlations. Cells with r^2^ > 0.4 are shaded by value and numerically annotated; all other cells are unshaded.

Pairwise correlations between per-sample process exposures were generally limited, with higher r^2^ values concentrated among selected groups (**Fig. 3b**). These included SBS12-associated processes, ultraviolet^9^-associated processes, MSI-associated^9,27^ processes containing SBS26/SBS44 and ID2, and APOBEC-associated^28–30^ processes containing SBS2 and SBS13. These patterns show that some separately inferred processes have related exposure distributions across samples. Their correspondence with established etiological groupings provides biological context for the inferred processes, although exposure correlation alone does not establish a shared mechanism.The exposure correlation structure also suggested finer groupings within these categories, motivating further examination of the UV-associated SBS7b and SBS12 variants described below.

### Joint reanalysis of PCAWG refines ultraviolet-associated mutational processes

Ultraviolet-associated mutagenesis^9,31,32^ provided an example of how joint multimodal inference can resolve variation within a canonical signature modality. MOTET identified an SBS7b-like signature within MP16 that differed from COSMIC SBS7b (cosine similarity = 0.79; **Fig. 4a**), with a lower contribution from T[C>T]C and a higher contribution from T[C>T]T.

**Fig. 4:**
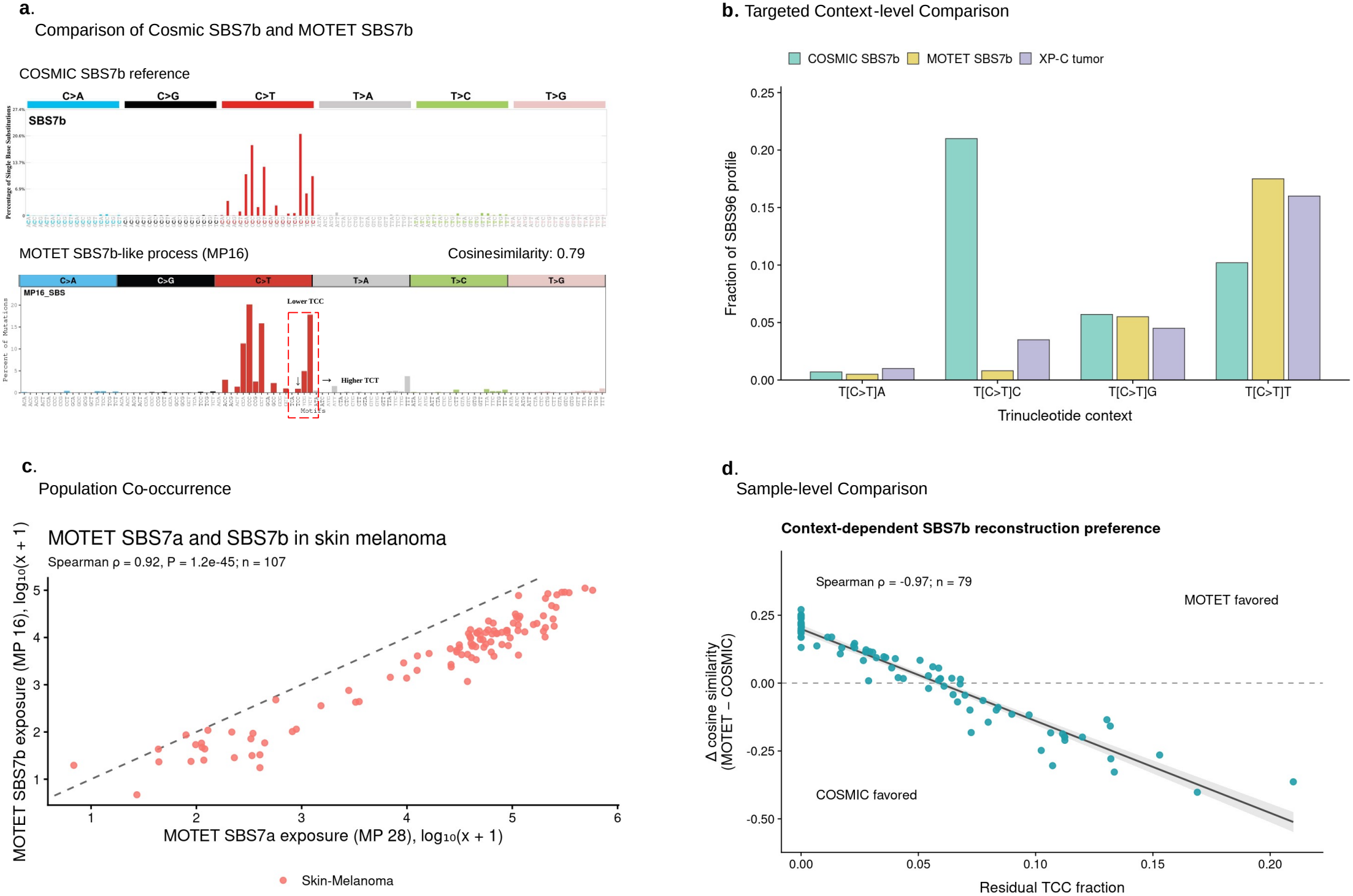
Joint reanalysis of PCAWG refines ultraviolet-associated mutational processes. **(a)** Comparison of the 96-channel COSMIC SBS7b profile (top) and the MOTET-derived SBS7b-like profile from MP16 (bottom; cosine similarity to COSMIC SBS7b = 0.79). The MOTET-derived profile has a lower contribution from T[C>T]C and a higher contribution from T[C>T]T relative to COSMIC SBS7b. **(b)** Comparison of T[C>T]N channels in COSMIC SBS7b, the MOTET-derived SBS7b-like profile, and the published XP-C skin cancer profile from Yurchenko et al (2023)^33^. Bar heights represent fractions of the respective SBS96 profiles. Both the MOTET-derived and XP-C profiles show lower T[C>T]C and higher T[C>T]T contributions than COSMIC SBS7b. **(c)** Association between per-sample exposures to the MOTET SBS7a-associated process (MP28) and SBS7b-like process (MP16) in PCAWG skin melanoma. Exposures are displayed after log_10_(x + 1) transformation. Each point represents one tumor; the dashed line indicates equal exposures. The Spearman correlation coefficient, two-sided P value, and sample size are indicated. **(d)** Relationship between the fraction of residual SBS mutations in the T[C>T]C channel and relative agreement with the two SBS7b profiles. Residual spectra were obtained by subtracting the fitted COSMIC SBS7a contribution from each tumor’s observed mutation spectrum. Analysis was restricted to 79 skin melanoma samples whose residual spectrum had cosine similarity ≥0.6 to either COSMIC SBS7b or the MOTET-derived SBS7b-like profile. Each point represents one tumor. The y-axis shows the difference in residual cosine similarity (MOTET-derived minus COSMIC SBS7b); positive values favor the MOTET-derived profile and negative values favor COSMIC SBS7b. The horizontal dashed line marks equal similarity. The solid line shows a linear regression fit, with shading indicating its 95% confidence interval. The Spearman correlation coefficient and sample size are indicated.

Comparison with the published XP-C skin cancer profile from Yurchenko et al. (2023)^33^ revealed a similar context shift: both the XP-C and MOTET-derived profiles showed lower T[C>T]C and higher T[C>T]T contributions than COSMIC SBS7b (**Fig. 4b**). This resemblance provides external support for the biological plausibility of the shifted profile. In addition, exposures to MP16 and the SBS7a-associated process MP28 (cosine similarity to COSMIC SBS7a = 0.986) were strongly correlated across PCAWG skin melanoma samples (Spearman ρ = 0.92, P = 1.2 × 10^−45^; **Fig. 4c**), consistent with their shared association with ultraviolet mutagenesis.

To examine how the MOTET-derived and COSMIC SBS7b profiles relate to sample-level variation, we compared their cosine similarities to residual spectra obtained after subtracting the fitted COSMIC SBS7a contribution from each tumour’s SBS96 spectrum (see Methods). We restricted this analysis to 79 melanoma samples whose residual spectrum had cosine similarity ≥0.6 to at least one of the two SBS7b profiles. Relative profile agreement, defined as the cosine similarity of the residual spectrum to the MOTET-derived profile minus that to COSMIC SBS7b, decreased with increasing residual T[C>T]C fraction (Spearman ρ = −0.97; **Fig. 4d**). Samples with higher residual T[C>T]C fractions therefore showed greater agreement with COSMIC SBS7b, consistent with the higher contribution of this channel to the reference profile.

At the process level, MOTET further distinguished the multimodal compositions of the SBS7a-associated process MP28 and the SBS7b-like process MP16. Both included DBS1-like components, whereas an ID13 component was assigned to MP28 but not MP16. Thus, despite their strongly correlated exposures, the two processes differed in their inferred indel associations. These findings suggest heterogeneity in the combinations of mutation modalities associated with ultraviolet mutagenesis, complementing the sample-level variation in SBS7b-like context composition.

### Joint reanalysis of PCAWG identifies putative SBS12 subtypes

MOTET identified two SBS12-related^2,9,30^ signatures within MP23 and MP18 (**Fig. 5a**). Their SBS components had cosine similarities of 0.75 and 0.86 to COSMIC SBS12, respectively, and their combined profile closely resembled the COSMIC reference (cosine similarity = 0.98; **Fig. 5b**). We provisionally designated these components SBS12a (MP23) and SBS12b (MP18) and examined their exposure patterns, reconstruction contributions and multimodal associations.

**Fig. 5:**
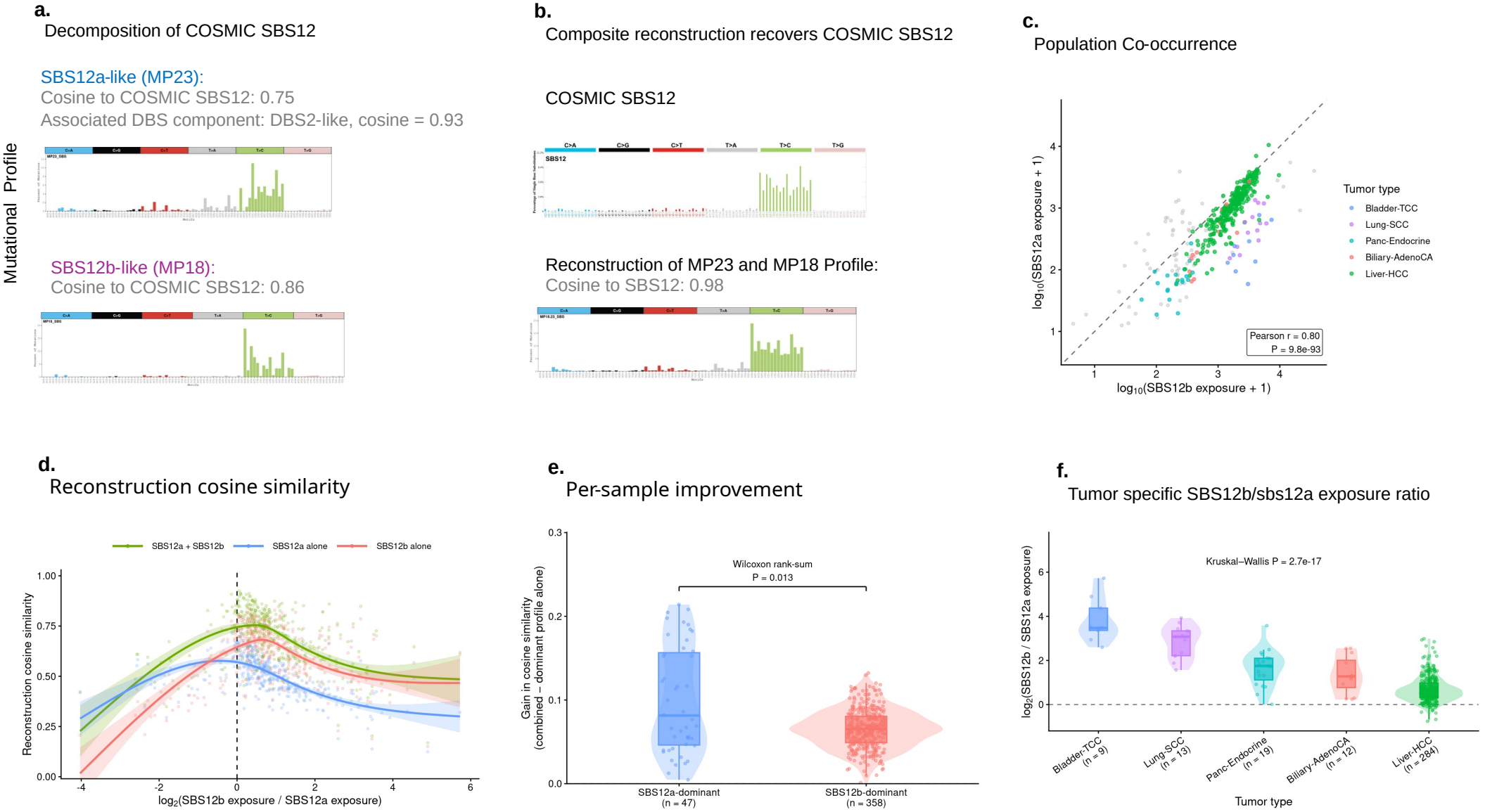
MOTET resolves two related SBS12-like components. **(a)** SBS12-like profiles inferred from the PCAWG cohort. MP23 contains an SBS12a-like profile associated with a DBS2-like component, whereas MP18 contains an SBS12b-like profile without a comparable multimodal association. The inferred SBS profiles have cosine similarities of 0.75 and 0.86, respectively, to COSMIC SBS12; the DBS component associated with MP23 has a cosine similarity of 0.93 to COSMIC DBS2. **(b)** Reconstruction of COSMIC SBS12 by combining the SBS12a-like MP23 and SBS12b-like MP18 profiles. The upper profile shows COSMIC SBS12 and the lower profile shows the combined reconstruction, which has a cosine similarity of 0.98 to COSMIC SBS12. **(c)** Per-sample exposures of SBS12a-like MP23 and SBS12b-like MP18 in 405 SBS12-active tumors, defined as tumors in which either process contributed >10 mutations or >7.5% of the total mutation burden, regardless of reconstruction similarity. Exposures are log_10_(x + 1)-transformed. Colors indicate the five tumor types with the most active samples; other types are gray. The dashed line marks equal exposure. Pearson correlation was calculated across all 405 tumors using the transformed exposures (r = 0.80; two-sided P = 9.8 × 10 ^−93^). **(d)** Cosine similarity of the observed SBS96 spectrum to reconstructions using SBS12a alone, SBS12b alone or both profiles in the same 405 tumors, plotted against the log_2_ exposure ratio (SBS12b/SBS12a). Negative values indicate SBS12a dominance; positive values indicate SBS12b dominance. The vertical dashed line marks equal exposure. Lines show generalized additive model fits with 95% confidence intervals. Similarities are calculated against the complete tumor spectrum, which also includes contributions from other signatures. **(e)** Per-sample gain in cosine similarity from combining SBS12a and SBS12b rather than using the exposure-dominant profile alone, compared between SBS12a-dominant (n = 47) and SBS12b-dominant (n = 358) tumors. Violins show kernel density estimates; boxes show medians and interquartile ranges, with whiskers extending to the most extreme values within 1.5 times the interquartile range. Points represent individual tumors. P = 0.013, two-sided Wilcoxon rank-sum test. **(f)**, Distribution of log_2_(SBS12b/SBS12a) exposure ratios across the five tumor types with the most SBS12-active tumors, ordered by decreasing median ratio. Sample sizes are shown below the labels; the horizontal dashed line marks equal exposure. Plot elements are defined as in e. Ratios differed among tumor types (Kruskal –Wallis test, χ^2^ = 83.8, d.f. = 4, P = 2.7 × 10^−17^).

The two components frequently co-occurred. Among 405 SBS12-active tumors, defined as tumors in which either component contributed >10 mutations or >7.5% of the total mutation burden, their log_10_(x + 1)-transformed exposures were strongly correlated (Pearson r = 0.80, P = 9.8 × 10^−93^; **Fig. 5c**). Across these tumors, reconstruction using both profiles generally showed greater cosine similarity to the observed SBS96 spectrum than reconstruction using either profile alone (**Fig. 5d)**. These comparisons used complete tumor spectra, which also contained contributions from other signatures. The gain over the exposure-dominant profile was greater in SBS12a-dominant tumors (n = 47) than in SBS12b-dominant tumors (n = 358; two-sided Wilcoxon rank-sum test, P = 0.013; **Fig. 5e**). Thus, inclusion of both profiles provided a larger improvement in SBS12a-dominant tumors.

The relative contributions of the two components also varied across tissues. Among the five tumor types with the most SBS12-active tumors, log_2_(SBS12b/SBS12a) exposure ratios differed substantially (Kruskal–Wallis χ^2^ = 83.8, d.f. = 4, P = 2.7 × 10^−17^; **Fig. 5f**). Median ratios were highest in Bladder-TCC and Lung-SCC, followed by Panc-Endocrine and Biliary-AdenoCA, while Liver-HCC showed the most balanced contributions. This variation indicates that the inferred components were not present in a fixed proportion across tissues, although it does not establish distinct etiologies.

The components also differed in their multimodal associations: MP23 contained the SBS12a-like component together with DBS2 and ID3, signatures associated with tobacco smoking^26^, whereas MP18 showed no comparable DBS/ID association. Together, the tissue-dependent exposure balance, differing reconstruction contributions and multimodal associations support SBS12a and SBS12b as candidate subtypes within the canonical SBS12 pattern. Further validation is needed to determine whether they reflect distinct underlying mechanisms.

### MOTET identifies a recurrent process linking an SBS37-related signature with a TG>CA-dominated DBS component

MOTET identified MP36 as a multimodal process containing an SBS37-related substitution signature and a DBS component most similar to COSMIC DBS17 (**Fig. 6a**). The SBS component had a cosine similarity of 0.845 to SBS37, supporting this assignment while leaving open whether the profile differences reflect incomplete separation from co-occurring signals or variation within the SBS37 pattern. MP36 activity was observed in Lymph-BNHL, Prost-AdenoCA, Panc-AdenoCA and Liver-HCC (Fig. 6d), tumor types in which SBS37 has also been reported^9,24^. This tissue overlap further supports the relationship between MP36’s substitution component and SBS37.

**Fig. 6:**
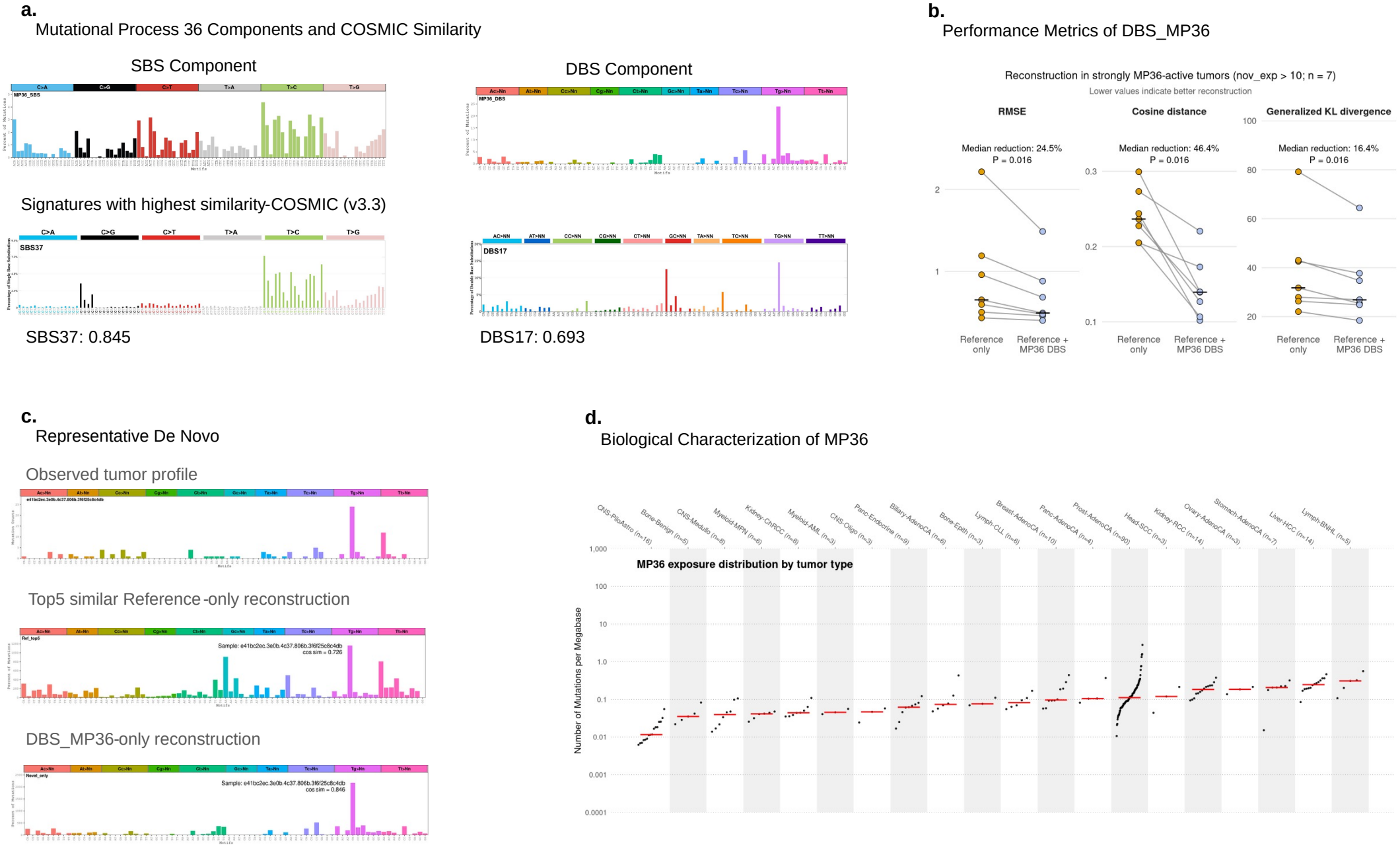
MOTET resolves SBS37-like signatures and novel DBS as components of a recurrent multimodal process. **(a)** SBS and DBS signature profiles inferred for MP36 (top), compared with their closest matches in COSMIC v3.6^24^, SBS37 and DBS17, respectively (bottom). Cosine similarities are shown below each profile pair. **(b)** DBS reconstruction accuracy in seven MP36-active tumors, defined by an inferred MP36 DBS exposure >10 mutations. Reconstructions use COSMIC DBS reference signatures alone (“Reference only”) or together with the MP36 DBS component (“Reference + MP36 DBS”). Accuracy is measured by per-sample root mean square error (RMSE), cosine distance and Kullback–Leibler (KL) divergence between observed and reconstructed spectra; lower values indicate better reconstruction. **(c)** DBS reconstruction for one representative tumor among the seven MP36-active tumors in b. Panels show the observed DBS profile (top), reconstruction using the five COSMIC DBS signatures with the highest cosine similarity to the MP36 DBS component (middle), and reconstruction using only the MP36 DBS component (bottom). (**d)** Distribution of MP36 exposure across tumor types in PCAWG_ALL, ordered by increasing median exposure. Each point represents one sample; red bars indicate tumor-type medians. Sample sizes are shown in parentheses.

The DBS component differed more substantially from its closest reference match, DBS17 (cosine similarity = 0.693; **Fig. 6a**). COSMIC describes DBS17 as containing prominent GC>AA and TG>CA contributions, with GC>AA also prominent in DBS^9,24^4. By contrast, the MP36 DBS profile was dominated by TG>CA, with a smaller GC>AA contribution. This pattern suggests that MP36 captures a more focused TG>CA-associated signal related to DBS17.

Adding the MP36 DBS component to the reference signature set improved reconstruction in seven tumors with inferred MP36 DBS exposure >10 mutations, reducing RMSE, cosine distance and KL divergence relative to reference signatures alone (Fig. 6b). A representative tumor illustrates reconstruction using the five COSMIC DBS signatures most similar to the MP36 component and, separately, using the MP36 component alone (Fig. 6c). These results support the utility of the inferred DBS profile for describing mutation spectra in MP36-active tumors, although they do not establish that the additional GC>AA contribution in COSMIC DBS17 represents noise.

In the SigProfiler^11^ analysis, the SBS component was only partially recovered, while the DBS signal was distributed across multiple components, including one resembling DBS4 more closely than DBS17. This comparison suggests that separate extraction can distribute related signals across components and leave their cross-modality relationships unresolved. By linking the SBS37-related and TG>CA-dominated DBS profiles within MP36, MOTET identifies a candidate cross-modality association whose underlying mechanism remains to be established.

## DISCUSSION

We developed MOTET, a Bayesian framework for jointly inferring mutational processes across single-base substitutions (SBSs), doublet-base substitutions (DBSs) and small insertions and deletions (IDs). Rather than decomposing each mutation modality independently and subsequently linking signatures through their exposures, MOTET models the three modalities as manifestations of a shared set of latent MPs. Applied to PCAWG whole-genome sequencing data, MOTET recovers processes consistent with well-established etiological associations, including the smoking-associated combination of SBS4, DBS2 and ID3^9,26^ and the ultraviolet-radiation-associated combination of SBS7, DBS1 and ID13^9^. These results demonstrate that joint inference can recover biologically coherent cross-modality relationships directly from mutation counts.

Joint modeling also suggests refinements to existing signature assignments. MOTET resolves SBS12-like activity into two processes with distinct tumor-type distributions and cross-modality characteristics (**Fig. 5**). Their profile similarities and overlap with the tumor-type distribution of SBS12 support their characterization as SBS12^9,30^-like components.

The analysis of MP36 illustrates how joint inference can connect incompletely characterized signatures across mutation modalities. MP36 links SBS37- and DBS17-like components within a shared process (**Fig. 6**), suggesting a potential shared origin for further investigation.

Joint extraction has previously been explored through concatenated mutation catalogs and shared-exposure models. PCAWG extracted composite signatures across 1,697 SBS, DBS and ID channels, recovering the tobacco-associated combination within a single composite signature^9^. Joint SBS and ID extraction also resolved homologous recombination deficiency signatures with different prognostic associations^16^. TensorSignatures links SNVs and other variants through a shared exposure matrix, with MNVs, indels and structural variants represented in an auxiliary block using custom classifications^15^. These studies establish precedents for using information across mutation modalities during signature discovery.

MOTET contributes to these joint approaches through a Bayesian hierarchical extension of MOTIVE that introduces a shared latent activity for each MP. Whereas MOTIVE models covariance among modality-specific activities, MOTET additionally represents cross-modality groupings explicitly, jointly estimating each process’s modality-specific profiles, allocation of mutations across modalities and activity in each tumor. MOTET recovers established COSMIC-annotated combinations and identifies additional candidate cross-modality associations, with these groupings inferred jointly from mutation counts rather than through post hoc exposure correlations (**Fig. 2**).

Several limitations should be considered. First, downsampling high-burden samples limited the influence of hypermutated tumors on signature discovery, a concern also addressed in previous pan-cancer analyses through separate extraction or count normalisation^9,11^, but reduced the mutation counts available to characterize them. More adaptive approaches to mutation-burden heterogeneity could better preserve this information while maintaining sensitivity across the broader cohort. Second, posterior sampling was computationally intensive, potentially limiting application to larger cohorts or expanded models. Improved parameterizations and approximate inference strategies, including variational^34^ or amortized^35^ inference, could improve scalability. Their effects on signature recovery and uncertainty calibration would need to be assessed against the current sampling-based approach. Finally, incomplete clinical annotation in PCAWG limited etiological characterization of newly identified processes and the ability to distinguish correlated demographic, environmental and treatment-related factors. Inferred cross-modality groupings do not by themselves establish a shared biological mechanism.

As whole-genome sequencing cohorts expand across tumor types, populations, treatment exposures and disease stages, joint inference may help resolve processes whose individual mutation-modality components are weak or ambiguous. Extending MOTET to incorporate copy-number alterations^36^ and structural variants^37^ could broaden its application to additional forms of somatic variation. Previous models have shown that point mutations and structural variants can be modelled jointly^15,21^. Together, these findings support MOTET as a framework for refining mutational signatures, characterizing their relationships across modalities and generating testable hypotheses about their biological origins.

## METHODS

### Model specification

#### Notation

Matrices are bold upper-case letters (e.g., **X**), vectors are bold lower-case letters (e.g., **x**), and scalars are plain lower-case letters (e.g., *x*). Count limits (numbers of samples, signatures, mutation types, etc.) are plain capital letters (e.g., *N, K, D*m). Subscripts index matrices and vectors using the corresponding lower-case letter (e.g., n indexes samples, so λn denotes exposures for sample *n*). Element-wise multiplication is denoted by ⊙. For an integer p ≥1), let Δ_p_ ⊂ ℝ^p^) denote the (p - 1)-dimensional probability simplex, Δ_p_ = {x∈ ℝ^p^ : *x*_j_ ≥ 0, 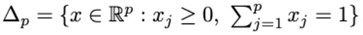 }. Table 1 lists the variables of MOTIF.

**Table 1:**
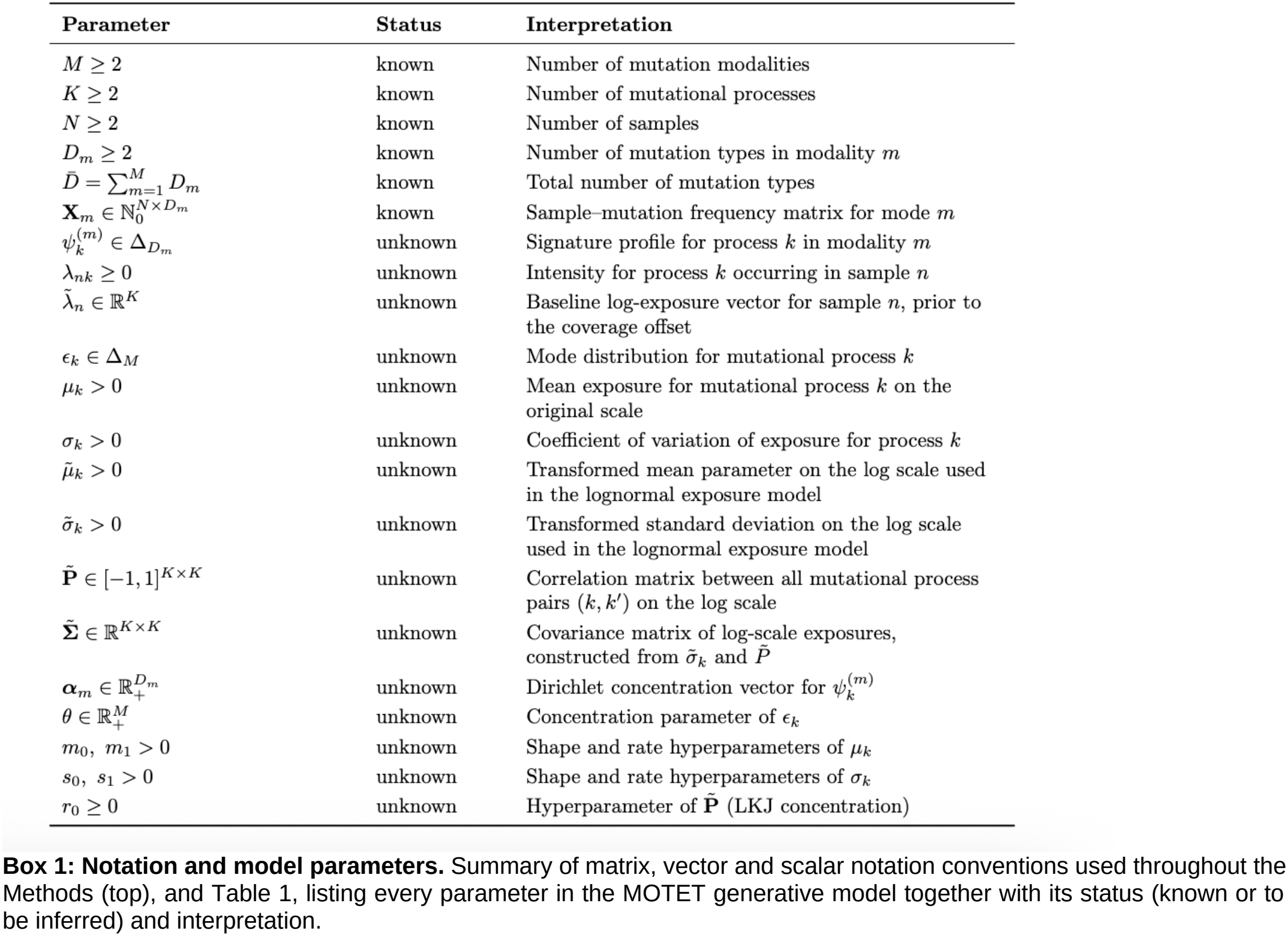
Parameters of MOTIF and their interpretation.

| Parameter | Status | Interpretation |
|---|---|---|
| M ≥ 2 | known | Number of mutation modalities |
| K ≥ 2 | known | Number of mutational processes |
| N ≥ 2 | known | Number of samples |
| D<sub>m</sub> ≥ 2 | known | Number of mutation types in modality m |
| $\bar{D} = \sum_{m=1}^{M} D_m$ | known | Total number of mutation types |
| $\mathbf{X}_m \in \mathbb{N}_0^{N \times D_m}$ | known | Sample–mutation frequency matrix for mode m |
| $\psi_k^{(m)} \in \Delta_{D_m}$ | unknown | Signature profile for process k in modality m |
| λ<sub>nk</sub> ≥ 0 | unknown | Intensity for process k occurring in sample n |
| λ̃<sub>n</sub> ∈ ℝ<sup>K</sup> | unknown | Baseline log-exposure vector for sample n, prior to the coverage offset |
| ϵ<sub>k</sub> ∈ Δ<sub>M</sub> | unknown | Mode distribution for mutational process k |
| μ<sub>k</sub> > 0 | unknown | Mean exposure for mutational process k on the original scale |
| σ<sub>k</sub> > 0 | unknown | Coefficient of variation of exposure for process k |
| μ̃<sub>k</sub> > 0 | unknown | Transformed mean parameter on the log scale used in the lognormal exposure model |
| σ̃<sub>k</sub> > 0 | unknown | Transformed standard deviation on the log scale used in the lognormal exposure model |
| **P̃** ∈ [−1, 1]<sup>K×K</sup> | unknown | Correlation matrix between all mutational process pairs (k, k′) on the log scale |
| **Σ̃** ∈ ℝ<sup>K×K</sup> | unknown | Covariance matrix of log-scale exposures, constructed from σ̃<sub>k</sub> and P̃ |
| $\boldsymbol{\alpha}_m \in \mathbb{R}_+^{D_m}$ | unknown | Dirichlet concentration vector for ψ<sub>k</sub><sup>(m)</sup> |
| θ ∈ ℝ<sub>+</sub><sup>M</sup> | unknown | Concentration parameter of ϵ<sub>k</sub> |
| m<sub>0</sub>, m<sub>1</sub> > 0 | unknown | Shape and rate hyperparameters of μ<sub>k</sub> |
| s<sub>0</sub>, s<sub>1</sub> > 0 | unknown | Shape and rate hyperparameters of σ<sub>k</sub> |
| r<sub>0</sub> ≥ 0 | unknown | Hyperparameter of **P̃** (LKJ concentration) |

We developed MOTET as a Bayesian framework for joint inference of multimodal mutational processes from SBS, DBS and ID mutation counts observed in the same samples, extending the MOTIVE model (Geyer, K. L. et al. Joint estimation of signatures across mutation modalities using multi-modal non-negative matrix factorization. Manuscript in preparation). Let 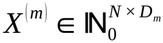 denote the mutation count matrix for modality *m∈* {1 ,*…, M* }, where *N* is the number of samples and *D*_*m*_is the number of mutation types in modality *m*. In this study, *M* = 3, corresponding to SBS, DBS and ID mutation modalities. MOTET assumes that observed mutation counts arise from *K* latent mutational processes shared across modalities.

For each process *k*, the model estimates a sample-level exposure *λ*_*nk*_, a modality distribution *ϵ*_*k*_ =(*ϵ*_*k* 1_ ,*…, ϵ*_*kM*_) *∈ Δ*_*M*_ (the (M−1)-dimensional probability simplex), and modality-specific mutational profiles 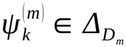 (the analogous simplex over the *D*_*m*_ mutation types in modality *m*). Here, *ϵ* _*km*_ denotes the relative contribution of process *k* to modality *m*, and 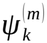 gives the relative rates of the *D*_*m*_ mutation types in modality *m*. This formulation provides a unified representation of mutational processes across mutation modalities and enables coordinated multimodal structure to be inferred within a single model.

For sample *n*, modality *m*, and mutation type *d*, the expected mutation count arises from three nested contributions. Each process *k* first contributes an expected *λ*_*nk*_ mutations to sample *n* in total, aggregated across all modalities and mutation types. The modality distribution *ϵ*_*k*_then partitions this budget across the *M* modalities: *λ*_*nk*_ *ϵ* _*km*_is the expected number of process-*k* mutations falling specifically in modality *m*, and since *ϵ*_*k*_ *∈ Δ*_*M*_, summing over *m* recovers the full exposure, 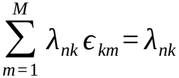. Within modality *m*, the profile *ψ*^(*m*)^ further partitions these *λ*_*nk*_ *ϵ*_*km*_ mutations across the *D* mutation types; since 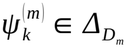 , summing over *d*returns *λ*_*nk*_ *ϵ*_*km*_ unchanged. Aggregating this per-process contribution over all *K* processes give the expected mutation count:

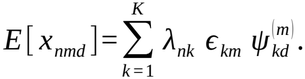

Observed counts are assumed to follow a Poisson distribution, as in Poisson non-negative matrix factorization^38,39^; and its Bayesian applications to mutational signatures^17,19^,

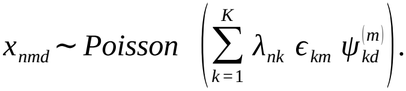

To account for residual variation in total mutation burden across samples, following the downsampling of high-burden outliers (see “Downsampling of high-burden samples”, below), we incorporated a sample-specific log-offset into the exposure model. For sample *n* and process *k*,

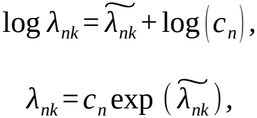

where *c*_*n*_ denotes the total mutation count for sample *n* after downsampling, summed across all M modalities, and 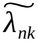 is the baseline log-exposure. This parameterization separates overall count magnitude from mutational pattern structure, so that expected counts scale proportionally with sample-level mutation burden while the latent process structure captures relative multimodal signature composition. This improves numerical stability during model fitting.

The baseline exposure vector for each sample was modeled on the log scale using a multivariate normal distribution, allowing correlated activities across processes^21^,

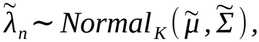

where 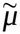 is the vector of transformed mean exposures and 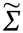 is the covariance matrix of log-scale exposures, constructed as 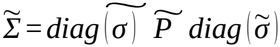^40^. The transformed parameters relate to the original-scale mean *μ*_*k*_ and coefficient of variation *σ*_*k*_ by standard lognormal moment-matching:

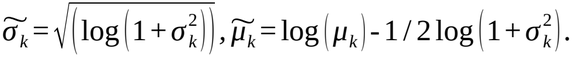

To allow processes with weak empirical support to be effectively pruned from an over-specified model, the original-scale mean exposure and coefficient of variation were themselves treated as unknown and assigned weakly informative priors, *μ*_*k*_ *∼ Inverse* - *Gamma*(*m*_0_ , *m*_1_) and *σ*_*k*_ *∼ Gamma*(*s*_0_ , *s*_1_), an automatic relevance determination (ARD)-style construction^17,41^ that shrinks the exposure of weakly supported processes toward zero (Algorithm 1).

The correlation matrix was assigned an LKJ prior^42^ 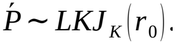*∼ LKJ* _*K*_ (*r*_0_) . Signature profiles were assigned modality-specific Dirichlet priors,

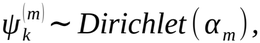

and modality distributions were assigned Dirichlet priors,

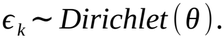

##### Algorithm 1 Generative process for MOTET with a sparsity-inducing ARD prior for mean exposure levels

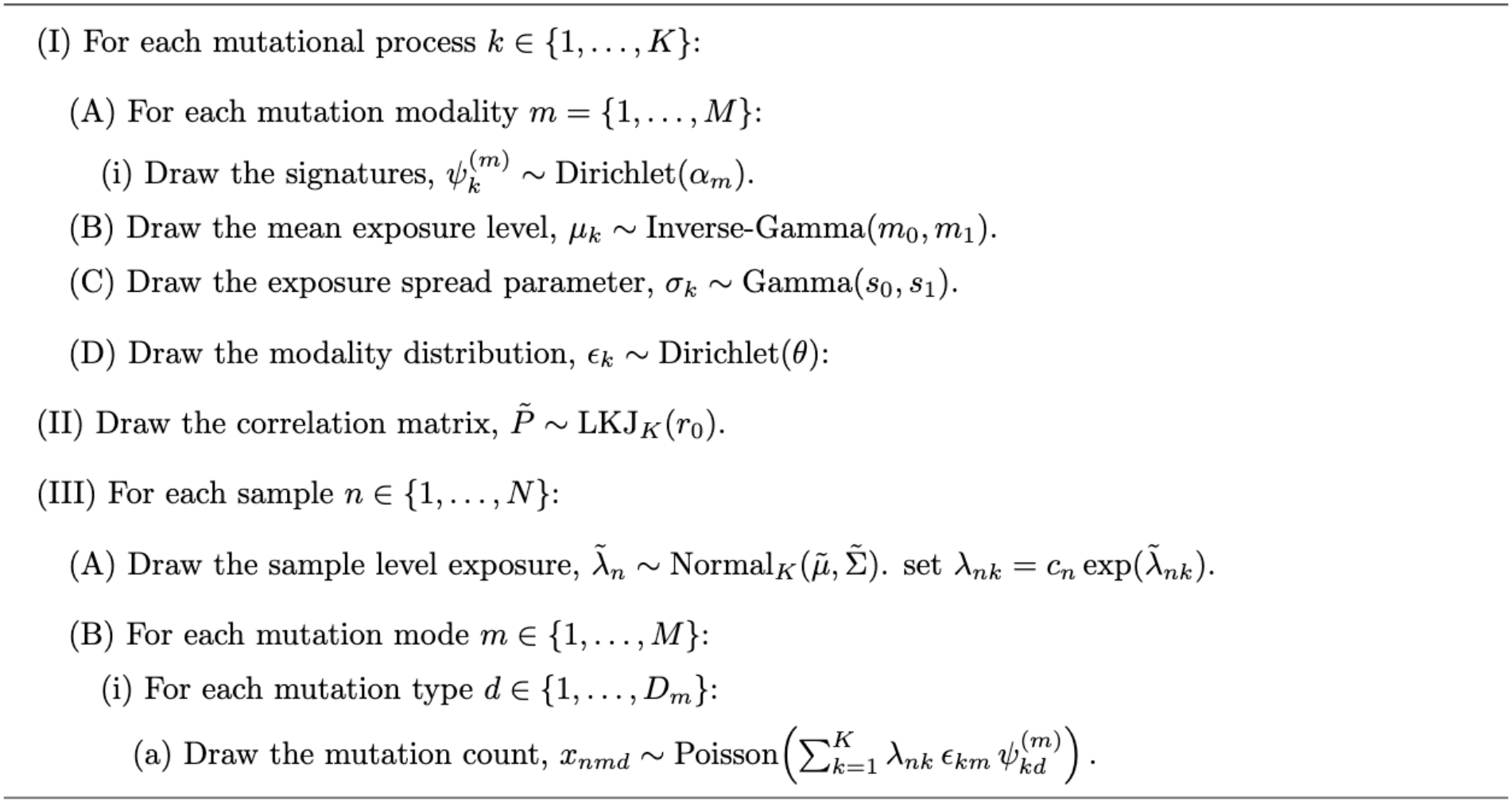

**Box 2: Generative process for MOTET**. Algorithm 1 specifies the full generative process underlying the model described above, including the priors on signature profiles, exposures and the process-correlation matrix, and the Poisson sampling distribution for observed mutation counts.

#### Model inference

Posterior inference for MOTET was performed in Stan^43^ (CmdStan v.2.36.0, via the R interface cmdstanr v.0.9.0^44^) using the No-U-Turn Sampler (NUTS)^45^, an adaptive Hamiltonian Monte Carlo algorithm. For each fitted model, we ran 10 independent chains for 20,000 iterations, comprising 15,000 warmup and 5,000 post-warmup iterations; warmup draws were discarded before posterior summarization. Sampling quality was assessed using effective sample size^46^, trace plots, and inspection of divergent transitions^47^, together with a within-chain split Rb computed separately for each chain by splitting it into two halves^46^. Convergence diagnostics were computed with the posterior R package (v.1.6.0)^48^.

Because between-chain heterogeneity increased with dataset size, we applied a chain-selection procedure to every fitted model, an approach motivated by the difficulty of mixing across modes in multimodal posteriors^49^. Chains were considered eligible for use if the 99th percentile of split Rb ^46^, fell below 1.05. For each eligible chain, we calculated (i) the mean log posterior density, as represented by Stan’s lp quantity^43^, and (ii) the Pritchard evidence heuristic, an approximation to the log marginal likelihood calculated from the mean and variance of the sampled log likelihoods^50^. Chains were ranked separately according to each statistic, and the two rankings were aggregated using a Borda count^51^. The chain with the highest aggregate rank was retained for downstream analysis.

### PCAWG cohort and preprocessing

#### Data acquisition

To enable a unified pan-cancer analysis, we assembled a PCAWG_ALL dataset comprising somatic mutation calls from 2,583 samples across 37 tumor types defined by the Pan-Cancer Analysis of Whole Genomes (PCAWG) Consortium^22^. We included only samples from donors on the PCAWG whitelist (Supplementary Table 1 of the original study^22^), which was defined using five quality-control metrics^52^. We use PCAWG histology abbreviations throughout. Supplementary Table 1 lists the corresponding full tumour type names and sample counts. We pooled variant calls (GRCh37/hg19) across tumour types and used musicatk (v2.0.0)^23^ to generate sample-level SBS96^2^, DBS78 and ID83^9^ mutation count matrices.

#### Downsampling of high-burden samples

In the PCAWG cohort, SBS counts reached approximately 2.4 million in a single sample. Samples with exceptionally high mutation burdens can dominate model fitting and reduce sensitivity to signatures with lower exposures^9^. Previous approaches have addressed this by extracting signatures separately in hypermutated samples^9^ or by normalizing per-sample counts before factorization^11^.

To limit the influence of high-burden samples on signature discovery, we downsampled mutation counts before fitting MOTET for the 259 of 2,583 PCAWG samples (10.0%) whose SBS counts exceeded the cohort’s 90th percentile (24,721.4 mutations). For each of these samples, we linearly mapped its SBS burden percentile from the 90th–100th percentile range to the 50th–90th percentile range, preserving the relative order of SBS burdens among adjusted samples. The corresponding empirical quantile of the cohort’s SBS counts gave the sample’s target SBS count. We multiplied the sample’s original DBS and ID totals by the ratio of its target to its original SBS count to obtain the respective target totals. Within each modality, we drew the target number of mutations by multinomial resampling, using that sample’s observed channel proportions as sampling probabilities.

For figures showing absolute MOTET exposures, we rescaled the fitted process exposures to the original mutation-count scale. For each downsampled sample, we multiplied all process exposures by the ratio of its original SBS count to its downsampled SBS count. All other samples had a scaling factor of 1.

#### Modality-specific sparsity priors for signature profiles

We specified separate Dirichlet concentration parameters for SBS, DBS and ID signature profiles (*α*_1_ = 0.5 , *α*_2_ = 0.25 , *α*_3_ = 0.25) to reflect differences in modality-specific signature structure. Because SBS signatures are generally denser across mutation contexts than DBS and ID signatures, a shared sparsity parameter (*α* = 1 for SBS, DBS and ID) tended to oversmooth the sparser DBS and ID profiles, yielding signatures that resembled mixtures of multiple underlying patterns. The modality-specific configuration improved separation and produced cleaner, more distinct DBS and ID signature estimates.

#### Classification and retention of inferred processes

Constituent signatures of each inferred process were compared with COSMIC Mutational Signatures v3.6 (GRCh37; https://cancer.sanger.ac.uk/signatures/)^24^ using cosine similarity in musicatk^23^, and were considered matched at a cosine similarity of ≥ 0.8. This threshold corresponds to the lower bound used in benchmarking of signature-extraction methods, below which similarity between non-negative profiles may arise by chance^11^. A process was classified as modality-dominant when its inferred proportion for SBS, DBS or ID exceeded 0.5, or, if no proportion exceeded 0.5, when COSMIC-matched signatures were present in only one modality. Processes without a qualifying COSMIC match were retained as candidate novel processes if they accounted for more than 40% of the total mutations in at least one sample, or more than 5% on average across the three samples with the highest proportional contributions. Processes without a qualifying COSMIC match that failed these criteria were classified as residual components.

#### Selecting the number of mutational processes

We used *K* = 40as an intentionally overcomplete specification for the PCAWG_ALL analysis. In a preliminary SigProfiler extraction at *K* = 40, 29 SBS components matched COSMIC reference signatures with cosine similarity *≥* 0.8. We allowed 11 additional components to accommodate MOTET’s joint modeling of SBS, DBS and ID mutations, which could recover modality-specific or novel processes not represented in the COSMIC SBS catalog. Thus, *K* = 40 was treated as a conservative upper bound rather than as the expected number of biological processes.

In the fitted *K* = 40 model, 38 of the 40 inferred components either contained a modality-specific profile matching a COSMIC v3.6 reference signature or met our criteria for retention as a novel process (see “Classification and retention of inferred processes”). The remaining two met neither criterion and were provisionally classified as unresolved, residual components. One retained component matched SBS111 signature^53^ (cosine similarity = 0.882), a signature recently added to COSMIC. Because SBS111 was not catalogued when the MOTET model was fitted, this post hoc match provides retrospective corroboration that the component captured structured biological signal rather than noise, although it does not constitute prospective validation.

### Comparison with separate extraction

#### SigProfilerExtractor analysis

For comparison with MOTET, we used SigProfilerExtractor (v1.1.24)^11^ to extract signatures separately from the original (non-downsampled) count matrix for each modality. We used its built-in Gaussian mixture model–based normalization to limit the influence of high-burden samples. For SBS96, we fixed the number of signatures at 40. For DBS78, we evaluated solutions with 5–10 signatures and retained the nine-signature solution recommended by SigProfilerExtractor. Extractions used 100 NMF replicates. The resulting de novo signatures were compared with COSMIC v3.6 reference signatures in musicatk^23^, as described above.

### Statistics and software

Analyses were performed in R (v.4.4.0)^54^ unless otherwise stated. MOTET was fitted in Stan^43^ (CmdStan v.2.36.0) via cmdstanr (v.0.9.0)^44^, and convergence diagnostics were computed with posterior (v.1.6.0)^48^. Mutation count matrices, COSMIC v3.6 signature comparisons and exposure estimation for the correlation baseline used musicatk (v.2.0.0)^23^. SigProfilerExtractor (v.1.1.24)^11^ was run in Python (v.3.10.15). Figures were generated with ggplot2 (v.4.0.0)^55^.

Similarity between signature profiles and mutation spectra was measured by cosine similarity. Associations between exposures were assessed by Pearson correlation r, as indicated in figure legends. For Fig. 5d, we fitted a separate generalized additive model to each reconstruction method’s cosine similarities using a shrinkage cubic regression spline (basis dimension k = 5) in mgcv (v.1.9.1)^56^, with the log_2_ ratio of SBS12b to SBS12a exposure as the predictor.

For Fig. 6b, we fitted each tumour’s DBS counts by non-negative least squares (nnls v.1.5)^57,58^ using reference DBS signatures with and without the MP36 DBS signature. We assessed reconstruction using root mean squared error across DBS channels, cosine distance and generalized Kullback–Leibler divergence from observed to reconstructed counts. Among tumours with MP36 DBS exposure >10, we compared each metric between fits using a two-sided paired Wilcoxon signed-rank test and reported unadjusted P values.

## Data availability

Somatic mutation calls from the Pan-Cancer Analysis of Whole Genomes (PCAWG) Consortium were obtained from https://www.synapse.org/Synapse:syn11726601/files/^22^. Reference mutational signatures were obtained from COSMIC Mutational Signatures v3.6 (GRCh37) https://cancer.sanger.ac.uk/signatures/.

Supplementary Table 1 lists the PCAWG tumour types analysed in this study. Supplementary Tables 2–4 provide the MOTET-inferred SBS, DBS and ID signature profiles, respectively. Supplementary Table 5 gives the proportion of mutations each process contributes to each modality. Supplementary Tables 6–8 provide per-sample exposures for SBS, DBS and ID, respectively.

## REFERENCES

1. Nik-Zainal, S. et al. Mutational processes molding the genomes of 21 breast cancers. Cell 149, 979–993 (2012).

2. Alexandrov, L. B. et al. Signatures of mutational processes in human cancer. Nature 500, 415–421 (2013).

3. Koh, G., Degasperi, A., Zou, X., Momen, S. & Nik-Zainal, S. Mutational signatures: emerging concepts, caveats and clinical applications. Nat. Rev. Cancer 21, 619–637 (2021).

4. Kucab, J. E. et al. A compendium of mutational signatures of environmental agents. Cell 177, 821-836.e16 (2019).

5. Pich, O. et al. The mutational footprints of cancer therapies. Nat. Genet. 51, 1732–1740 (2019).

6. Davies, H. et al. HRDetect is a predictor of BRCA1 and BRCA2 deficiency based on mutational signatures. Nat. Med. 23, 517–525 (2017).

7. Gerstung, M. et al. The evolutionary history of 2,658 cancers. Nature 578, 122–128 (2020).

8. Tate, J. G. et al. COSMIC: the Catalogue Of Somatic Mutations In Cancer. Nucleic Acids Res. 47, D941– D947 (2019).

9. Alexandrov, L. B. et al. The repertoire of mutational signatures in human cancer. Nature 578, 94–101 (2020).

10. Alexandrov, L. B., Nik-Zainal, S., Wedge, D. C., Campbell, P. J. & Stratton, M. R. Deciphering signatures of mutational processes operative in human cancer. Cell Rep. 3, 246–259 (2013).

11. Islam, S. M. A. et al. Uncovering novel mutational signatures by de novo extraction with SigProfilerExtractor. Cell Genomics 2, 100179 (2022).

12. Degasperi, A. et al. Substitution mutational signatures in whole-genome–sequenced cancers in the UK population. Science 376, abl9283 (2022).

13. Jin, H. et al. Accurate and sensitive mutational signature analysis with MuSiCal. Nat. Genet. 56, 541–552 (2024).

14. Gruber, A. J. et al. Integrated signatures define mutational processes in prostate cancer. Nature https://doi.org/10.1038/s41586-026-10468-w (2026) doi:10.1038/s41586-026-10468-w.

15. Vöhringer, H., van Hoeck, A., Cuppen, E. & Gerstung, M. Learning mutational signatures and their multidimensional genomic properties with TensorSignatures. Nat. Commun. 12, 3628 (2021).

16. Ferrer-Torres, P., Galván-Femenía, I. & Supek, F. Joint inference of mutational signatures from indels and single-nucleotide substitutions reveals prognostic impact of DNA repair deficiencies. Genome Med. 17, 76 (2025).

17. Kim, J. et al. Somatic ERCC2 mutations are associated with a distinct genomic signature in urothelial tumors. Nat. Genet. 48, 600–606 (2016).

18. Grabski, I. N., Trippa, L. & Parmigiani, G. Bayesian multi-study non-negative matrix factorization for mutational signatures. Genome Biol. 26, 98 (2025).

19. Xue, C., Miller, J. W., Carter, S. L. & Huggins, J. H. Robust discovery of mutational signatures using power posteriors. PLOS Comput. Biol. 22, e1014372 (2026).

20. Buscaroli, E. et al. BASCULE: Bayesian inference and clustering of mutational signatures leveraging biological priors. Genome Biol. 27, 15 (2025).

21. Funnell, T. et al. Integrated structural variation and point mutation signatures in cancer genomes using correlated topic models. PLOS Comput. Biol. 15, e1006799 (2019).

22. The ICGC/TCGA Pan-Cancer Analysis of Whole Genomes Consortium et al. Pan-cancer analysis of whole genomes. Nature 578, 82–93 (2020).

23. Chevalier, A. et al. The Mutational Signature Comprehensive Analysis Toolkit (musicatk) for the Discovery, Prediction, and Exploration of Mutational Signatures. Cancer Res. 81, 5813–5817 (2021).

24. Sondka, Z. et al. COSMIC: a curated database of somatic variants and clinical data for cancer. Nucleic Acids Res. 52, D1210–D1217 (2024).

25. Nik-Zainal, S. et al. Landscape of somatic mutations in 560 breast cancer whole-genome sequences. Nature 534, 47–54 (2016).

26. Alexandrov, L. B. et al. Mutational signatures associated with tobacco smoking in human cancer. Science 354, 618–622 (2016).

27. Meier, B. et al. Mutational signatures of DNA mismatch repair deficiency in C. elegans and human cancers. Genome Res. 28, 666–675 (2018).

28. Roberts, S. A. et al. An APOBEC cytidine deaminase mutagenesis pattern is widespread in human cancers. Nat. Genet. 45, 970–976 (2013).

29. Petljak, M. et al. Mechanisms of APOBEC3 mutagenesis in human cancer cells. Nature 607, 799–807 (2022).

30. Letouzé, E. et al. Mutational signatures reveal the dynamic interplay of risk factors and cellular processes during liver tumorigenesis. Nat. Commun. 8, 1315 (2017).

31. Brash, D. E. UV Signature Mutations. Photochem. Photobiol. 91, 15–26 (2015).

32. Hayward, N. K. et al. Whole-genome landscapes of major melanoma subtypes. Nature 545, 175–180 (2017).

33. Yurchenko, A. A. et al. Genomic mutation landscape of skin cancers from DNA repair-deficient xeroderma pigmentosum patients. Nat. Commun. 14, 2561 (2023).

34. Blei, D. M., Kucukelbir, A. & McAuliffe, J. D. Variational Inference: A Review for Statisticians. J. Am. Stat. Assoc. 112, 859–877 (2017).

35. Kingma, D. P. & Welling, M. Auto-Encoding Variational Bayes. In 2nd International Conference on Learning Representations (ICLR 2014) (arXiv, Banff, Canada, 2014).

36. Steele, C. D. et al. Signatures of copy number alterations in human cancer. Nature 606, 984–991 (2022).

37. Li, Y. et al. Patterns of somatic structural variation in human cancer genomes. Nature 578, 112–121 (2020).

38. Lee, D. D. & Seung, H. S. Learning the parts of objects by non-negative matrix factorization. Nature 401, 788–791 (1999).

39. Cemgil, A. T. Bayesian Inference for Nonnegative Matrix Factorisation Models. Comput. Intell. Neurosci. 2009, 785152 (2009).

40. Barnard, J., McCulloch, R. & Meng, X.-L. Modeling covariance matrices in terms of standard deviations and correlations, with application to shrinkage. Stat. Sin. 10, 1281–1311 (2000).

41. Tan, V. Y. F. & Fevotte, C. Automatic Relevance Determination in Nonnegative Matrix Factorization with the /spl beta/-Divergence. IEEE Trans. Pattern Anal. Mach. Intell. 35, 1592–1605 (2013).

42. Lewandowski, D., Kurowicka, D. & Joe, H. Generating random correlation matrices based on vines and extended onion method. J. Multivar. Anal. 100, 1989–2001 (2009).

43. Carpenter, B. et al. Stan : A Probabilistic Programming Language. J. Stat. Softw. 76, (2017).

44. Gabry, J., Češnovar, R., Johnson, A. & Bronder, S. cmdstanr: R Interface to ‘CmdStan’. GitHub (2025).

45. Hoffman, M. D. & Gelman, A. The No-U-Turn Sampler: adaptively setting path lengths in Hamiltonian Monte Carlo. J Mach Learn Res 15, 1593–1623 (2014).

46. Vehtari, A., Gelman, A., Simpson, D., Carpenter, B. & Bürkner, P.-C. Rank-Normalization, Folding, and Localization: An Improved R̂ for Assessing Convergence of MCMC (with Discussion). Bayesian Anal. 16, 667–718 (2021).

47. Betancourt, M. A Conceptual Introduction to Hamiltonian Monte Carlo. Preprint at 10.48550/arXiv.1701.02434 (2018).

48. Bürkner, P.-C., Gabry, J., Kay, M. & Vehtari, A. posterior: Tools for Working with Posterior Distributions. 1.7.0 10.32614/CRAN.package.posterior (2021).

49. Yao, Y., Vehtari, A. & Gelman, A. Stacking for non-mixing Bayesian computations: the curse and blessing of multimodal posteriors. J. Mach. Learn. Res. 23, 1–45 (2022).

50. Pritchard, J. K., Stephens, M. & Donnelly, P. Inference of Population Structure Using Multilocus Genotype Data. Genetics 155, 945–959 (2000).

51. Dwork, C., Kumar, R., Naor, M. & Sivakumar, D. Rank aggregation methods for the Web. In Proceedings of the 10th international conference on World Wide Web 613–622 (ACM, Hong Kong Hong Kong, 2001). doi:10.1145/371920.372165.

52. Whalley, J. P. et al. Framework for quality assessment of whole genome cancer sequences. Nat. Commun. 11, 5040 (2020).

53. Kazachkova, M. et al. Identification and Validation of a Previously Missed Mutational Signature in Colorectal Cancer. Preprint at 10.1101/2025.06.06.658321 (2025).

54. R Core Team. R: A Language and Environment for Statistical Computing. R Foundation for Statistical Computing (2024).

55. Wickham, H. Ggplot2: Elegant Graphics for Data Analysis. (Springer International Publishing : Imprint: Springer, Cham, 2016). doi:10.1007/978-3-319-24277-4.

56. Wood, S. N. Fast Stable Restricted Maximum Likelihood and Marginal Likelihood Estimation of Semiparametric Generalized Linear Models. J. R. Stat. Soc. Ser. B Stat. Methodol. 73, 3–36 (2011).

57. Lawson, C. L. & Hanson, R. J. Solving Least Squares Problems. (Society for Industrial and Applied Mathematics, Philadelphia, 1995). doi:10.1137/1.9781611971217.

58. Mullen, K. M. & van Stokkum, I. H. M. nnls: The Lawson-Hanson Algorithm for Non-Negative Least Squares (NNLS). CRAN (2023).

